# Decoding Promoter Activity from DNA Sequence using Pre-trained Language Models

**DOI:** 10.64898/2026.02.05.704015

**Authors:** Christophe Jung

**Affiliations:** Gene Center and Department of Biochemistry, Quantitative and Molecular Biology (QMB), Ludwig-Maximilians-Universität München, Butenandtstr. 1, 81377 München, Germany

## Abstract

Promoter architecture plays a central role in transcriptional regulation, yet predicting promoter activity directly from DNA sequence remains challenging. Here, we assess whether transformer-based DNA language models can learn and interpret regulatory logic encoded in *Drosophila* core promoters. We fine-tuned the 117-million-parameter DNABERT-2 model on ∼700 synthetic promoters assayed by ∼2,600 dual-luciferase measurements in *Drosophila* S2 cells. A sequence-only model achieved high predictive accuracy (R² ≈ 0.91), demonstrating that core promoter sequence alone strongly constrains transcriptional output. Model interpretability using SHapley Additive exPlanations (SHAP) revealed biologically meaningful sequence features corresponding to canonical core promoter elements. Extending the model to incorporate biological context, including Ecdysone (Ecd) hormonal activation and flanking −1/+1 nucleosomal sequences, preserved strong performance while capturing more complex promoter behavior. Gene-wise cross-validation showed robust generalization for most promoters, and application to independent *in vivo* embryo data demonstrated that the model still generalizes reasonably well, even in complex biological contexts.

## INTRODUCTION

Accurate control of gene expression is fundamental to development and cellular function.^1,2^ In metazoans, transcription initiation is largely determined by the core promoter, a region of approximately 150 bp surrounding the transcription start site (TSS) that recruits RNA polymerase II and the general transcription machinery.^3,4^ Core promoters integrate several types of regulatory information, including short sequence motifs,^5,6^ motif strength and spacing,^7–9^ DNA shape,^10^ and local chromatin context.^11–13^ In *Drosophila melanogaster*, systematic experimental work has identified a diverse set of core promoter elements (CPEs), such as the Initiator (INR),^5^ TATA-box,^14^ downstream promoter element (DPE/MTEDPE),^4,15^ Ohler motifs,^7,16^ and others, which combine in distinct architectures associated with developmental or constitutive gene expression.^13^

Massively parallel reporter assays (MPRAs)^17,18^ have enabled direct and quantitative testing of promoter sequences at large scale. These studies showed that promoter activity can often be explained by additive effects of individual motifs, with additional influence by motif position,^18^ sequence context,^18^ nucleosome positioning,^16^ and hormonal signaling.^13^

Despite these advances, building predictive models that generalize across promoters and genes remains difficult. In our previous work,^13^ promoter activity in *Drosophila* S2 cells was successfully modelled using linear regression with motif-based features, demonstrating that expression can be largely explained by additive contributions of individual motifs. However, such models depend on predefined motif annotations and do not test whether promoter regulation can be learned directly from DNA sequence.

Transformer-based DNA language models offer powerful alternative for modelling regulatory DNA.^19,20^ DNABERT-2^20^ adapts the BERT (Bidirectional Encoder Representations from Transformers)^21^ architecture to genomic sequences by learning contextualized representations of overlapping k-mers through masked-language pretraining. This allows the model to capture short sequence motifs as well as their positional and combinatorial relationships without motif predefinitions.

Sequence-based language models therefore provide a way to test whether core promoter activity can be predicted from DNA sequence alone, while also offering the possibility of biological interpretation. While such models have shown promise for tasks such as chromatin accessibility and enhancer prediction,^22^ their ability to quantitatively predict promoter activity and to yield biologically interpretable insights has not been systematically tested.^22^

Here, we fine-tune DNABERT-2 on a large synthetic promoter dataset that we previously generated,^13^ which contains quantitative measurements of *Drosophila* core promoter activity under controlled experimental conditions. We evaluate predictive performance and use model interpretability methods to identify which sequence features drive promoter activity predictions. We further integrate biological context and test whether the model generalizes across genes and on independent *in vivo* data. Together, our results show that DNA language models can accurately predict promoter activity from sequence and recover known principles of core promoter regulation.

Fig. 1 provides an overview of the experimental and computational framework used in this study. Synthetic core promoter variants were assayed in *Drosophila* S2 cells using a quantitative dual-luciferase reporter system,^13^ yielding reproducible measurements of promoter activity with and without Ecd hormonal stimulation (Fig. 1A,B).

**Figure 1.**
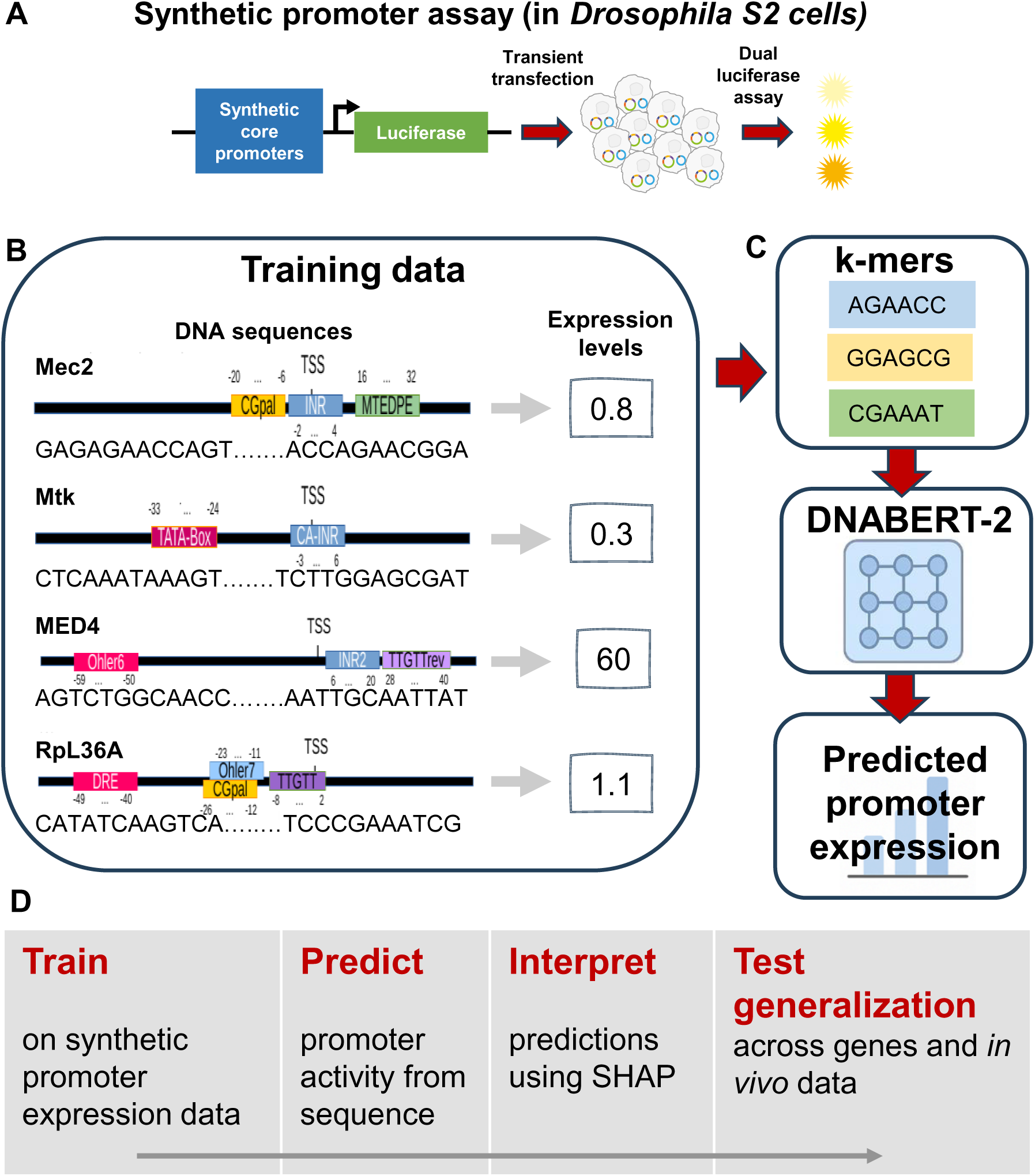
Experimental system and modeling overview. **(A)** Schematic of the dual-luciferase reporter assay used to quantify synthetic core promoter activity in *Drosophila* S2 cells (data from Qi et al.^13^). **(B)** Training dataset consisting of designed promoter DNA sequences paired with experimentally measured expression levels, used for model fine-tuning. **(C)** Schematic depicting the k-merisation process of the input DNA sequence, that will be used by the DNABERT-2 algorithm for promoter expression predictions. **(D)** Computational workflow: promoter sequences are used to train a sequence-based model, predictions are interpreted using SHAP, biological context is integrated, and model generalization is assessed using gene-wise validation and independent *in vivo* data.

These sequence-expression pairs were used to fine-tune an adapted DNABERT-2 model to predict promoter activity directly from DNA sequence (Fig. 1C). Model predictions were evaluated on held-out data and validated using control experiments. To interpret the learned regulatory logic, we used SHAP^23^ to assign contributions of individual sequence features to predicted expression levels. Biological context, including Ecd activation and flanking −1/+1 nucleosomal sequences, was also incorporated to examine how sequence-intrinsic promoter logic is modulated by regulatory environment. Finally, we assessed model generalization using gene-wise cross-validation and independent *in vivo* promoter activity data from *Drosophila* embryos (Fig. 1D).

## RESULTS

### Fine-tuning DNABERT-2 on synthetic Drosophila promoter libraries

We fine-tuned DNABERT-2 on a subset of our previously generated synthetic promoter dataset,^13^ focusing first on core promoter variants measured in a fixed nucleosomal context (referred to N1.x and N7.y, corresponding to −1 and +1 nucleosomal sequences,^13^ respectively), and without Ecd induction. The dataset comprised approximately 700 unique promoter sequences (131 bp core region) with a total of ∼2,600 quantitative luciferase measurements, averaging ∼2.8 biological replicates per sequence. Expression values were log2-transformed prior to modelling.

DNABERT-2 supports native subword-based tokenization schemes that generate many short and variable-length sequence tokens.^20^ While these representations yielded comparable predictive performance in preliminary experiments (data not shown), they strongly reduced interpretability because sequence features were split across many tokens. In contrast, fixed 6-mer tokenization provided a simpler and more biologically intuitive representation, aligned with known core promoter motifs and enabled robust interpretation using SHAP. We therefore adopted 6-mer tokenization for all analyses (Methods).

We randomly split individual promoter activity measurements into 90% training and 10% held-out test sets. The model was fine-tuned for regression using mean squared error (MSE) loss.

The fine-tuned sequence-only model achieved strong predictive performance on the held-out test set (R² ≈ 0.91; MSE ≈ 0.7), indicating close agreement between predicted and measured promoter activity across a wide dynamic range (Fig. 2A and Supplementary Fig 1A). Residuals were symmetrically distributed with no strong systematic bias across expression levels (Fig. 2B).

**Figure 2.**
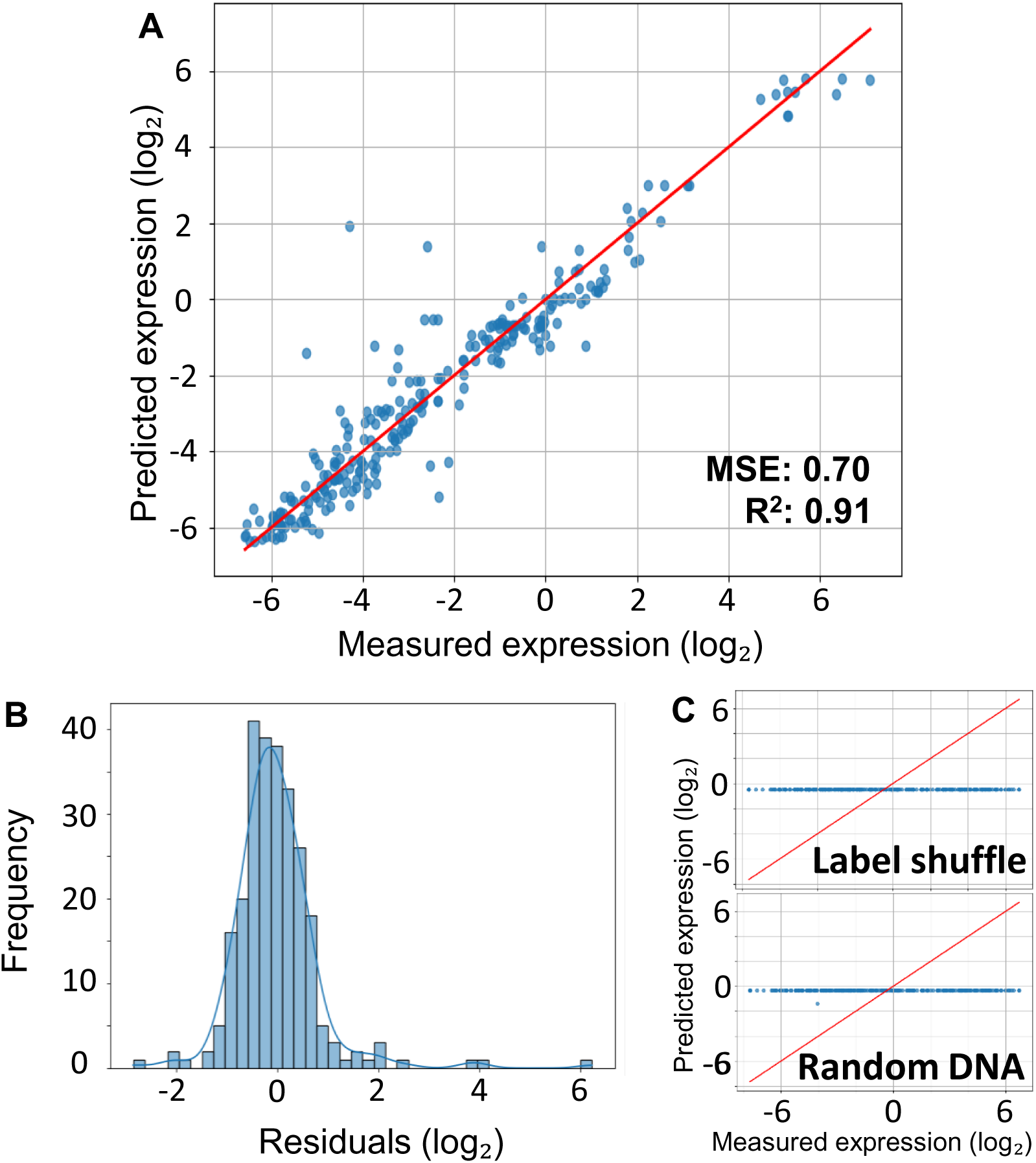
Accurate prediction of promoter activity from DNA sequence. **(A)** Scatter plot comparing predicted and measured promoter activity (log₂ scale) for promoter sequences randomly held out from training using a standard train/test split. The red diagonal indicates perfect prediction. **(B)** Distribution of prediction residuals across the activity range. **(C)** Negative controls. Predicted versus measured activity after shuffling expression labels (upper panel) or replacing promoter sequences with random DNA (lower panel). In both cases, predictive performance collapses.

To verify that the model really learned sequence-expression relationships, we performed negative control experiments. When expression labels were randomly shuffled across sequences, or when promoter sequences were replaced by random DNA of matched length and composition (Methods), predictive performance collapsed to baseline levels (Fig. 2C upper and lower panels, respectively and Supplementary Fig. 1BC). These controls demonstrate that DNABERT-2 relies on real regulatory sequence information rather than memorization or trivial correlations.

### Model interpretability reveals promoter regulatory logic

Transformer-based models learn attention weights that quantify how strongly individual sequence tokens attend to one another when making predictions.^19,20^ In principle, attention maps can highlight regions of a DNA sequence that the model considers informative. However, for promoter activity prediction, attention maps alone are difficult to interpret biologically. DNABERT-2 contains many layers and attention heads, resulting in complex patterns that cannot be easily linked to specific regulatory elements (Supplementary Fig. 2).

Therefore, we used SHAP to obtain direct and quantitative insight into which sequence features drive predictions. This method assigns an additive contribution value to each input token based on its effect on the predicted output.^23^ This allows promoter activation or repression to be directly attributed to specific sequence elements, making SHAP particularly well suited for promoter analysis.

To quantify how each input affects the prediction, we applied SHAP to the fine-tuned model (Methods), assigning importance scores to each 6-mer based on its contribution to predicted expression (Fig. 3). Across the dataset, SHAP consistently highlighted short sequence elements matching known core promoter motifs. SHAP-enriched 6mers were associated with known core promoter motifs by comparison to *XXmotif* consensus sequences,^7,13^ allowing for limited sequence variation (Methods).

**Figure 3.**
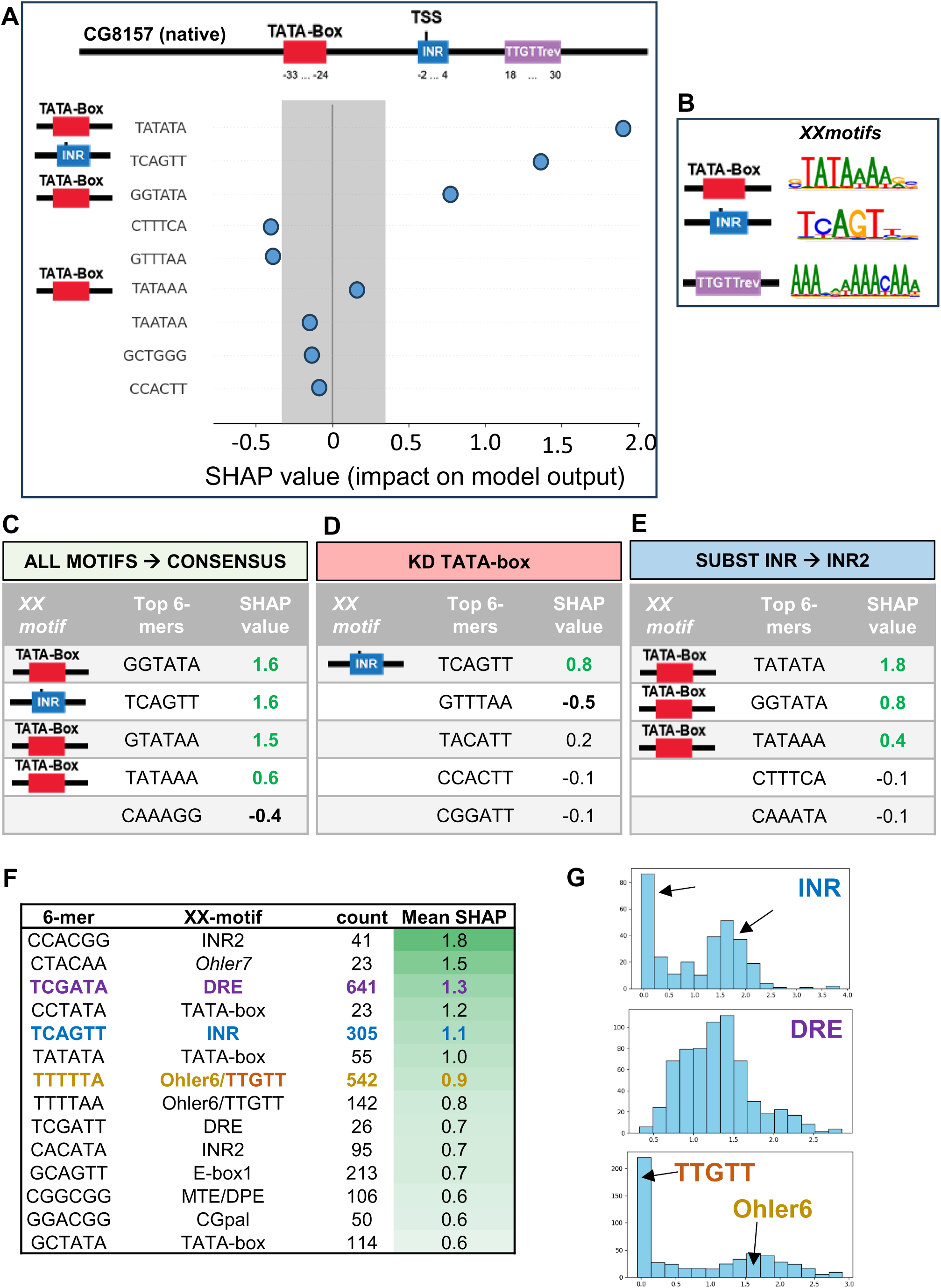
SHAP analysis reveals promoter regulatory logic learned by the model. **(A)** SHAP attribution scores mapped along the sequence of a representative promoter (native CG8157). SHAP-enriched 6-mers were assigned to motif classes (on the left hand-side) based on sequence similarity to *XXmotif* consensus motifs. The upper panel indicate the promoter architecture with motif positions as found by *XXmotif*.^13^ **(B)** Binding preferences for the three motifs identified by *XXmotif* in CG8157. **(C-E)** SHAP attribution profiles for promoter variants carrying defined motif perturbations. Shown are representative cases for CG8157 including **C r**eplacement of all motifs by their consensus sequences, **D** motif knockout of the TATA-box, and **E** motif substitution (Initiator replaced by INR2). Compared to the native promoter profile shown in **A**, replacement by consensus motifs results in stronger and more focused SHAP signals, disruption of the TATA-box leads to a loss of SHAP signal at the corresponding positions, while functional substitution redistributes SHAP contributions and INR2 does not recover the loss of INR. **(F)** Table of the most informative 6-mers across the dataset, ranked by mean SHAP contribution. For each 6-mer, the associated *XXmotif* annotation, occurrence count, and mean SHAP value are shown. Highly ranked 6-mers correspond to canonical core promoter elements. The colors used to highlight SHAP-enriched 6-mers match the ones used for the motif-specific SHAP value distributions shown in **G**. **(G)** Distributions of SHAP values for representative motif classes. Initiator-associated 6-mers show a bimodal distribution (upper panel), DRE-associated 6-mers display a broad distribution (midlle panel), and TTGTT/Ohler6-associated 6-mers show overlapping bimodal distributions, reflecting differences in positional constraints and regulatory roles (lower panel).

For the developmental promoter CG8157, SHAP-enriched regions showed strong agreement with experimentally defined promoter elements identified by de novo motif discovery (Fig. 3A,B). INR and TATA box motifs contributed strong positive effects, whereas the TTGTTrev motif did not appear among the top SHAP values, consistent with its previously reported weakly repressive role.

Systematic motif perturbations revealed that SHAP attributions responded in a specific and predictable way to changes in motif identity and strength. Replacing native motifs with consensus sequences (Fig. 3C) led to amplified and more sharply localized positive contributions, indicating increased promoter strength. Knockdown of the TATA box (Fig. 3D) caused a near-complete loss of SHAP signal at the TATA position and reduced positive contributions across the promoter, with no compensatory effects from other motifs. Substitution of the Initiator motif (Fig. 3E) with an alternative variant (INR2) redistributed SHAP signal to the new motif context while preserving partial activity. Interestingly, INR2 did not recover INR SHAP signal.

A global analysis of the most informative 6-mers further supported the conclusion that the highest-impact tokens corresponded to well-known core promoter motifs, including INR, INR2, TATA box, DRE, Ohler6/7, and MTEDPE (Fig. 3F). Differences in SHAP value distributions across motif classes reflected known positional constraints and regulatory roles (Fig. 3G–I). Together, these results demonstrate that DNABERT-2 learns biologically meaningful promoter grammar and that SHAP provides a powerful framework for interpreting learned sequence features.

### Integrating biological context modulates prediction accuracy

While core promoter sequence defines a strong baseline for expression, biological context such as hormonal signaling and chromatin environment modulates transcriptional output. As established previously,^13^ developmental and constitutive promoters differ markedly in their responsiveness to hormonal activation, while nucleosomal sequences flanking the core promoter exert additional, though more modest, effects on expression. These context dependencies are summarized in Fig. 4A-C. Developmental promoters show significantly higher ecdysone inducibility than constitutive promoters (Fig. 4A-B; Wilcoxon rank-sum test, p = 0.0054), whereas −1/+1 nucleosomal sequences modulate expression levels more moderately, and in a promoter dependent manner (Fig. 4C).

**Figure 4.**
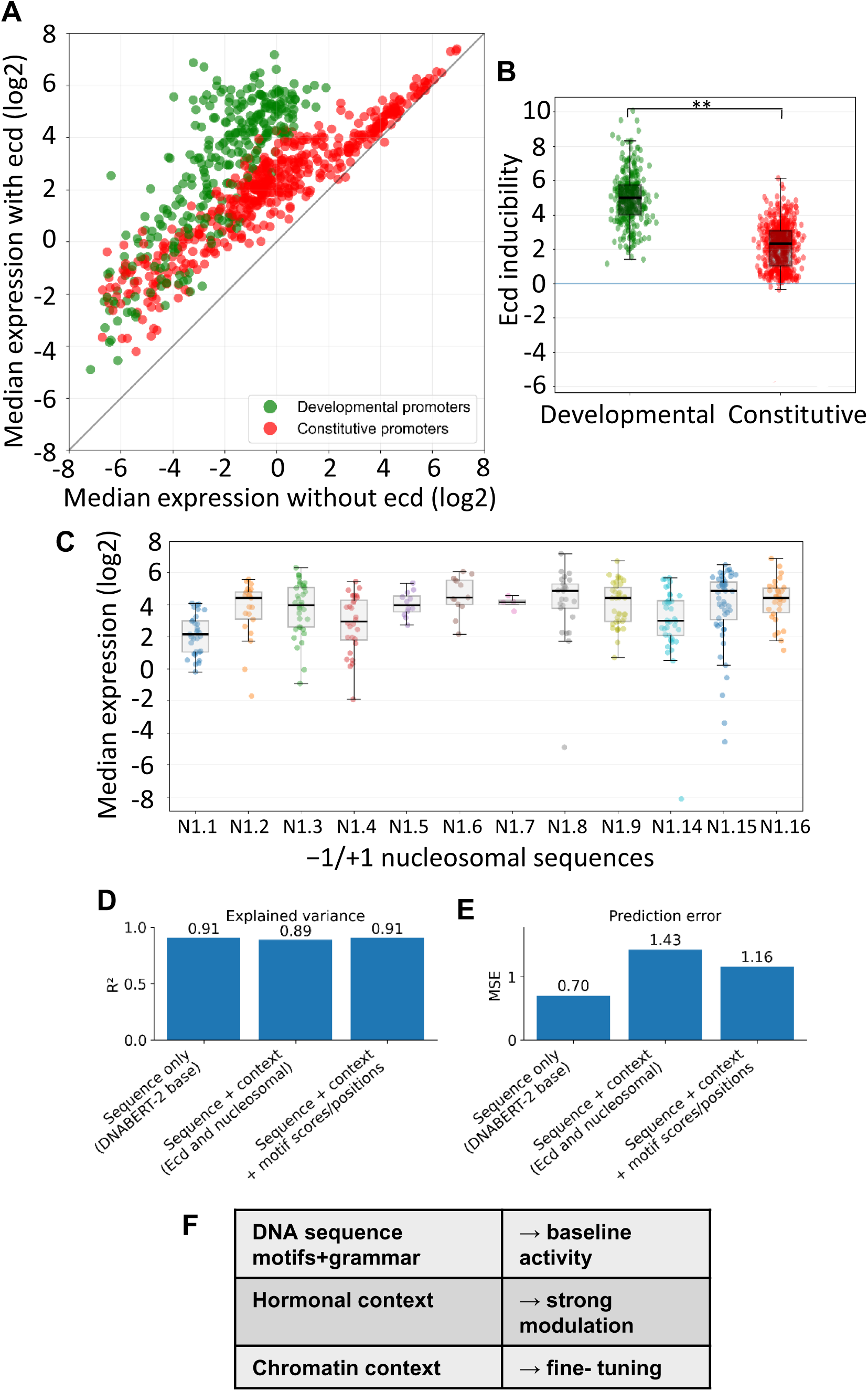
Biological context modulates promoter activity and prediction accuracy. **(A)** Hormonal induction of promoter activity. Median promoter activity measured in the presence of Ecd (log₂ scale) plotted against median activity measured without Ecd (log₂ scale) for all our synthetic promoter variants. Each point represents one promoter. The diagonal indicates equal expression in both conditions. Data are derived from Qi et al.^13^ **(B)** Ecdysone inducibility (Δlog₂ expression) grouped by promoter class. As previously reported,^13^ developmental promoters show significantly higher inducibility than constitutive promoters (p = 0.0054). Statistical significance was assessed using a two-sided Wilcoxon rank-sum test. **\*** p < 0.05; **\*\*** p < 0.01; **\*\*\*** p < 0.001; ns, not significant. Boxes indicate median and interquartile range; points represent individual promoters. **(C)** Effect of −1/+1 nucleosomal sequences on promoter activity. Median promoter activity (log₂ scale) plotted for different −1/+1 nucleosomal sequences flanking the core promoter (N1.y variants). Each point represents the median activity of promoter variants measured in the same nucleosomal context. Data are derived from Qi et al.^13^ **(D-E)** Effect of biological context on prediction accuracy. Model performance is shown for sequence-only input and for models incorporating biological context. **D** Explained variance (R²) and **E** prediction error (MSE) are shown. While the sequence-only model already achieves high performance (R² ≈ 0.91), inclusion of ecdysone and −1/+1 nucleosomal sequences maintains strong performance (R² ≈ 0.89), and adding explicit motif features leads to a modest increase in explained variance. **(F)** Conceptual summary of promoter regulation. DNA sequence, including motif grammar, defines baseline promoter activity, which is strongly modulated by hormonal input and fine-tuned by −1/+1 nucleosomal sequences.

To test whether these effects can be integrated with sequence-based promoter logic, we extended the DNABERT-based framework to jointly model promoter sequence and biological context. In addition to the core promoter sequence, the model incorporates the Ecd activation state and quantitative features describing the −1 and +1 nucleosomal flanking sequences. These features were concatenated with the sequence embedding and passed through additional fully connected layers before regression (Methods).

Including biological context preserved strong predictive performance while capturing known regulatory effects (Fig. 4D,E). The model correctly reproduced the higher ecdysone responsiveness of developmental promoters, as well as the weaker but consistent influence of nucleosomal sequences on expression. Adding explicit motif-derived features provided only marginal improvement (MSE: 1.43 to 1.16; R²: 0.89 to 0.91), indicating that most information related to motifs is already encoded in the sequence representation (Fig. 4F).

Together, these results show that DNA language models can be naturally extended to integrate heterogeneous biological context, yielding a unified framework in which promoter sequence defines baseline activity and the biological context signals modulate transcription in a predictable and biologically meaningful manner.

### Gene-wise cross-validation reveals promoter class-specific generalization

A key question is whether the model can generalize to promoters from genes that were not seen during training. To address this, we performed gene-wise cross-validation, in which all promoter variants from a given gene were held out as the validation set and the model was trained from scratch on promoters from all remaining genes. This procedure prevents information leakage and directly tests generalization across genes.^18^

Figure 5A–C shows representative examples of gene-wise held-out predictions for promoters with different architectures. For the developmental promoter cas, predicted and measured activities showed good agreement (MSE = 2.31, R² = 0.81; Fig. 5A). Similarly strong performance was observed for the constitutive promoter THR (MSE = 0.93, R² = 0.89; Fig. 5B). In contrast, predictions for the motif-less promoter CG15674 were more variable (MSE = 2.44, R² = 0.40; Fig. 5C), illustrating that gene-wise performance can differ between individual promoters.

**Figure 5.**
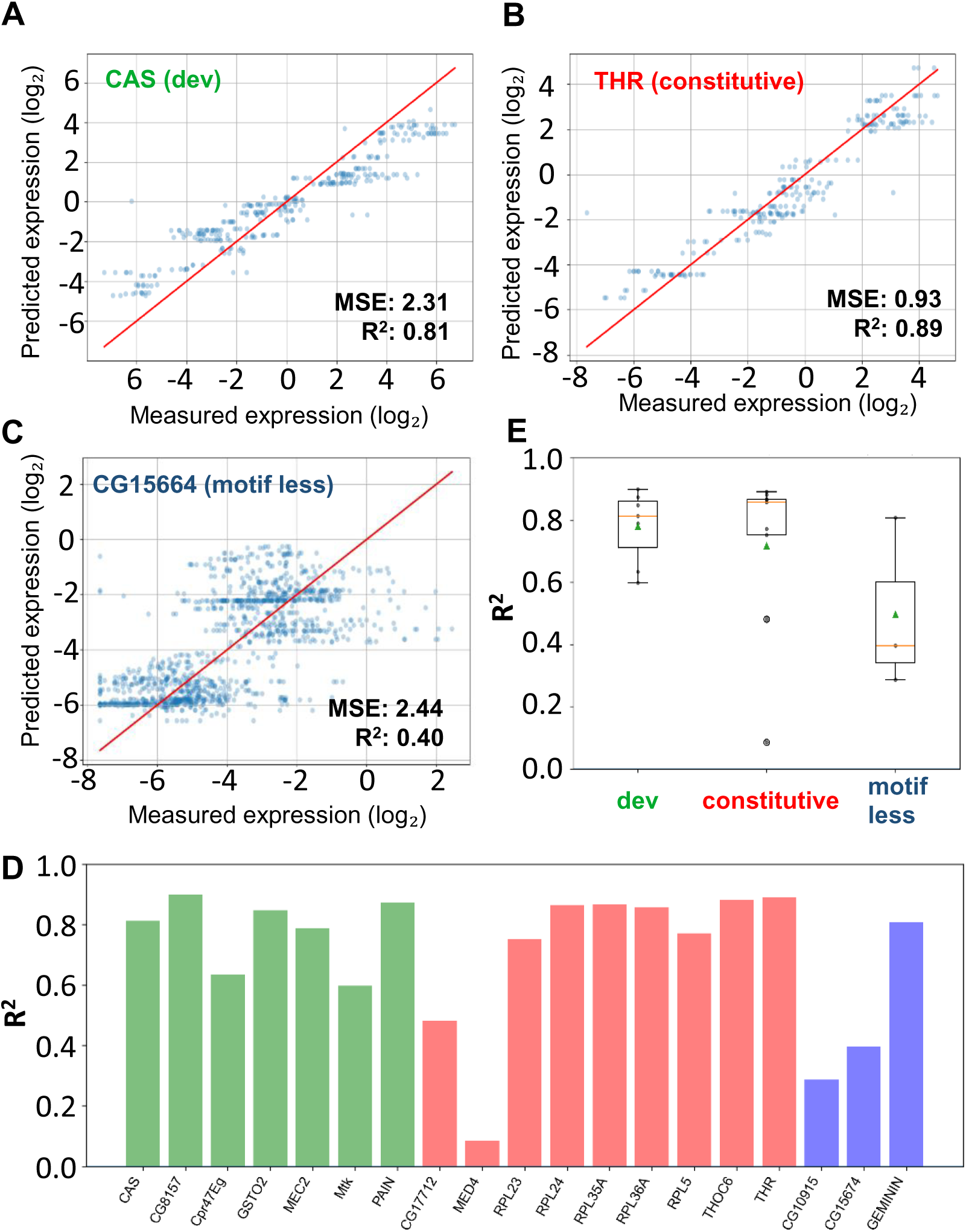
Gene-wise held-out validation reveals class-specific generalization. **(A)** Representative examples of held-out genes. Predicted versus measured promoter activity is shown for representative developmental, constitutive, and motif-less promoters, illustrating successful and failed generalization at the level of promoter variants. **(B)** Distribution of gene-wise prediction performance (R²) grouped by promoter class. Boxes indicate median and interquartile range; whiskers show the full data range. No significant differences were observed between promoter classes (one-way ANOVA, F = 1.67, p = 0.22). **(C)** Gene-wise prediction performance when all promoter variants from a given gene are excluded from training. Each point represents one gene and is coloured by promoter class.

Across genes, gene-wise prediction performance was generally high (Fig. 5D), with most promoters showing substantial explained variance between predicted and measured activity. Developmental promoters were consistently well predicted (R² typically ∼0.6-0.9; Fig. 5D-E), while many constitutive promoters, particularly ribosomal and housekeeping genes such as RpL24, RpL35A, RpL36A, Thoc6, and THR, also exhibited strong generalization (R² ≈ 0.85-0.90). A small number of promoters, including MED4 and CG17712, showed reduced performance. Motif-less promoters showed more variable performance, with lower R² values for CG10915 and CG15674, but strong performance for GEMININ (R² = 0.81).

When grouped by promoter class, developmental and constitutive promoters showed slightly higher median prediction performance than motif-less promoters (Fig. 5E). However, these differences were modest and not statistically significant (one-way ANOVA, F = 1.67, p = 0.22), indicating that generalization depends more on individual gene properties than on promoter class alone.

### Generalization to *in vivo* Drosophila embryo promoter activity

Finally, to test generalization beyond the synthetic *in vitro* system, we evaluated our model on an independent *in vivo* dataset from Li et al.,^24^ which measured the activity of randomly mutated developmental promoters (*svb* and DSCP) in *Drosophila* embryos (Fig. 6A). This dataset provides a challenging benchmark because transcription *in vivo* occurs in a complex biological environment.

**Figure 6.**
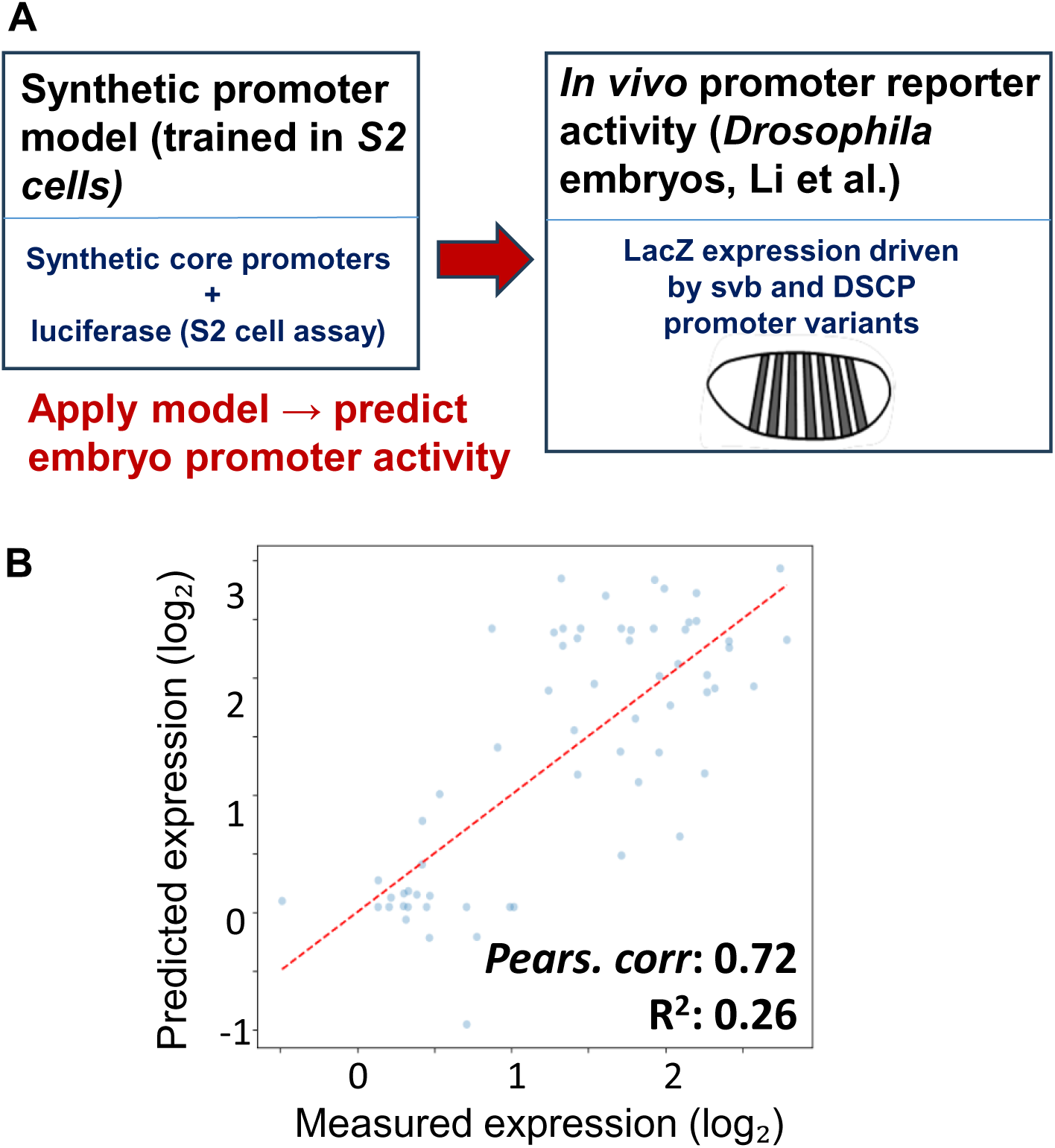
Generalization to in vivo *Drosophila* embryo promoter activity. **(A)** Schematic illustrating the validation strategy of model performance on an independent *Drosophila* embryo dataset. A model trained exclusively on synthetic promoter activity measured in S2 cells is applied without retraining to promoter sequences from an independent *in vivo* embryo reporter assay.^24^ **(B)** Scatter plot showing Pearson correlation and coefficient of determination (R²) between promoter activity predicted by our model and measured activity in *Drosophila* embryos.

Without retraining or parameter adjustment, the model achieved moderate predictive performance (Pearson correlation of r ≈ 0.63 and an R² ≈ 0.21) and captured major trends in promoter activity (Fig. 6B). Performance accuracy was lower than in S2 cells, which is expected given the influence of enhancer-promoter interactions,^22^ chromatin accessibility,^22^ and developmental timing *in vivo*,^18^ factors that are not encoded in the core promoter sequence.

Together, these results indicate that core promoter sequence explains a substantial, but incomplete, component of transcriptional regulation *in vivo*.

## DISCUSSION

Our study demonstrates that DNA language models can accurately predict quantitative promoter activity directly from sequence and provide biologically interpretable insights into promoter regulation. Fine-tuned DNABERT-2 explained over 90% of the variance in synthetic promoter expression, extending traditional motif-based models.

Interpretability analyses showed that the model recovers canonical core promoter elements without explicit motif annotation. SHAP analysis revealed clear contributions from INR, TATA box, DPE/MTEDPE, and Ohler motifs, consistent with prior experimental studies. Differences in SHAP value distributions across motif classes reflected known regulatory roles and positional preferences.

Integrating biological context preserved strong predictive performance while capturing known modulatory effects of hormonal signalling and nucleosomal environment. Genewise cross-validation demonstrated robust generalization for most developmental and constitutive promoters, whereas performance for motif-less promoters was more heterogeneous. This behaviour is consistent with the known dependence of motif-less promoters on chromatin state and higher-order regulatory mechanisms.

Validation on *in vivo* embryo data showed moderate generalization, indicating that core promoter sequence alone cannot fully explain gene expression in complex developmental contexts. This is expected, because transcription *in vivo* depends on additional regulatory layers, including enhancer-promoter interactions, chromatin accessibility, transcription factor availability, and developmental timing.

A main strength of our approach is that it combines quantitative promoter activity prediction with clear interpretability, allowing regulatory sequence features to be identified directly from DNA without predefined motif annotations. In contrast to recent deep learning models for regulatory genomics, such as DeepSEA,^25^ Basenji,^26^ and Enformer,^22^ which focus on genome-wide prediction accuracy, our model is trained on controlled synthetic promoter measurements, making biological interpretation of learned features more straightforward using SHAP.

The method also has important limitations. It is trained on synthetic promoter libraries in a single cellular context and uses a sequence-only, fixed-length promoter window, which cannot capture long-range regulation or cell-type specific chromatin states. As a result, the approach is best suited for studying core promoter logic rather than fully predicting gene expression in complex biological systems. Future models that include chromatin or other regulatory information may help overcome these limitations.

Overall, our results establish transformer-based DNA language models as accurate and interpretable tools for studying promoter regulation. Beyond basic understanding, these models provide a quantitative framework for rational promoter design and synthetic biology applications.

## METHODS

### Synthetic promoter dataset

We used the synthetic promoter dataset generated in our previous work,^13^ which systematically perturbs *Drosophila* core promoter sequences and measures their activity using a dual-luciferase reporter assay in S2 cells.^27^ Each construct consists of a −1 nucleosomal sequence (N1.x), an enhancer module, a 131 bp core promoter region centered on the transcription start site (TSS), and a +1 nucleosomal sequence (N7.y). Promoter activity was quantified as Firefly/Renilla luciferase ratios and normalized across experiments.

For the base sequence-only model, we restricted the dataset to promoter variants measured without ecdysone induction and flanked by fixed −1 and +1 nucleosomal sequences (N1.x and N7.y)^13^ Expression values were log₂-transformed after clipping low values (minimum 5×10⁻⁴) to ensure numerical stability. Unless otherwise stated, individual biological replicates were treated as independent measurements during model training.

### Sequence preprocessing and k-mer tokenization

Promoter sequences were trimmed to the 131 bp core promoter region. Sequences were tokenized using overlapping fixed-length k-mers (k = 6, stride = 1), yielding 126 k-mer tokens per sequence. Following the DNABERT-2 input convention, special [CLS] and [SEP] tokens were added.

Although DNABERT-2 natively supports subword-based tokenization, we empirically found that fixed 6-mer tokenization provided comparable predictive performance while enabling substantially improved interpretability. Six-mers closely match the length scale of canonical core promoter motifs and allowed stable attribution of sequence features using SHAP.

### Model architecture and fine-tuning

We fine-tuned the pre-trained DNABERT-2 (117M parameters) transformer model for regression by replacing the classification head with a single linear output neuron. Model weights were initialized from the publicly available DNABERT-2 checkpoint.

Models were trained using mean squared error (MSE) loss and optimized with *AdamW*. Unless otherwise noted, hyperparameters were: learning rate 1×10⁻⁵, batch size 28, weight decay 4–6×10⁻⁴, and 25 training epochs. Dropout probabilities in both hidden and attention layers were set to very low values (≈1×10⁻⁵) to stabilize regression. Training was performed on NVIDIA GPUs (GeForce GTX 1080 or equivalent). Training typically required 30–120 minutes per model.

### Train–test splitting and gene-wise cross-validation strategies

For standard evaluation, promoter measurements were randomly split into 90% training and 10% held-out test sets. To assess generalization beyond individual genes, we additionally performed gene-wise cross-validation, in which all promoter variants derived from a given gene were held out entirely as the test set, and a new model was trained from scratch using promoter variants from all other genes. This process was repeated for each gene, ensuring strict separation between training and validation data.

Model performance was evaluated using mean squared error (MSE), coefficient of determination (R²), and Pearson correlation coefficient.

### Context-aware models

To incorporate biological context beyond core promoter sequence, we extended the base DNABERT-2 model to include additional inputs: (i) a binary indicator of Ecd induction, and (ii) quantitative features representing the −1 and +1 nucleosomal sequences (N1.x and N7.y^13^). These features were concatenated with the DNABERT-derived sequence representation and processed through two additional fully connected layers prior to regression.

In a further extension, explicit motif-derived features were included. These comprised PWM-based motif scores and positional information for a curated set of known *Drosophila* core promoter motifs. Motif features were standardized using z-score normalization based on the training set only. While this model yielded marginal performance gains, the increased complexity led us to use the sequence + context model as the primary context-aware architecture.

### Model interpretability and feature attribution

To interpret sequence-based predictions, we applied SHapley Additive exPlanations (SHAP) at the k-mer level. SHAP values were computed using a background distribution of randomly generated DNA sequences matched in length to the input promoters. For each sequence, SHAP attribution scores were assigned to individual 6-mer tokens, indicating their contribution to predicted expression.

SHAP values were aggregated across sequences to identify globally informative k-mers and to compare their distributions across promoter classes. For selected promoters, SHAP scores were visualized along the sequence and compared with independently identified motif annotations.

To associate SHAP-enriched 6-mers with known core promoter elements, we compared high-impact 6-mers to consensus motifs identified previously using *XXmotif*.^13^ This mapping was performed manually by sequence comparison, allowing up to two single-nucleotide mismatches relative to the *XXmotif*-derived consensus sequences. This approach enabled approximate assignment of SHAP-enriched k-mers to canonical promoter motifs while accounting for motif degeneracy and local sequence variability

### Attention analysis

To complement SHAP-based interpretation, attention weights were extracted from all transformer layers and heads during inference. Attention maps were visualized for individual promoters and averaged across layers, heads, and sequences to identify consistently attended regions. As attention patterns were highly distributed and complex, these analyses were used qualitatively and reported in Supplementary Fig. 2.

### Prediction for the *In vivo* validation

External validation used published *in vivo* embryo reporter data from Li et al.,^24^ measuring the activity of mutated *svb* and DSCP promoters. The DNABERT-2 model trained on S2 cell data was applied directly to these sequences without retraining. To ensure compatibility with the DNABERT-2 model, we aligned the transcription start site (TSS) of each Li et al. sequence to the TSS and extracted a 131 bp window centered on the TSS, matching the core promoter length used throughout this study.

The DNABERT-2 model trained on S2 cell data was then applied directly to these cropped sequences without retraining or parameter tuning. Predicted promoter activities were compared to the corresponding measured expression values to evaluate generalization across experimental systems.

## SOFTWARE AND REPRODUCIBILITY

All analyses were performed in *Python* using *PyTorch* and the *HuggingFace* Transformers library. Custom scripts were used for model training, prediction, interpretability analyses, and visualization. Model configurations, training parameters, and evaluation outputs were logged and exported to enable reproducibility.

## DATA AVAILIBILITY

All experimental datasets analyzed in this study have been published previously. Synthetic promoter activity data measured in *Drosophila* S2 cells are available from Qi *et al.* (2022).^13^ In vivo embryo promoter activity data were obtained from Li *et al.* (2023).^24^

## CODE AVAILIBILITY

All custom code used for model training, evaluation, and interpretability analyses will be made publicly available on GitHub upon publication.

## ACKNOWLEDGMENTS

We are grateful to Xueying Li and Justin Crocker for kindly providing access to the full *in vivo* Drosophila embryo promoter dataset used for external validation in this study This work was supported by the Gene Center Munich and the Graduate School for Quantitative and Molecular Biology (QMB), Ludwig-Maximilians-Universität München.

## AUTHOR CONTRIBUTIONS

C.J. conceived the study, performed all computational analyses, interpreted the results, and wrote the manuscript.

## AI USAGE DISCLOSURE STATEMENT

Custom Python scripts were developed by the author. AI-assisted tools (ChatGPT, OpenAI; GitHub Copilot, Microsoft) were used during manuscript drafting for language editing, clarity, and iterative text refinement. All analyses, code, data interpretation, and scientific conclusions were performed, reviewed, and validated by the author. AI tools were not used for data generation, analysis or interpretation.

## SUPPLEMENTARY FIGURES

**Supplementary Figure 1.** Model training stability and controls Validation loss curves.

**(A)** for the base sequence-only model.

**(B)** when expression labels were randomly shuffled across sequences (Methods).

**(C)** when promoter sequences were replaced by random DNA of matched length and composition (Methods).

**Supplementary Figure 2.** Attention map visualizations

**(A)** Representative attention maps from the transformer model. DNABERT-2 contains 12 transformer layers with 12 attention heads each, yielding 144 attention matrices per sequence, each of size ∼130×130. While averaged attention highlighted regions near the TSS (Token position ∼80) and downstream elements, the complexity of these patterns limited direct biological interpretation. The heatmap color scale (right) indicates normalized attention weights, with higher values corresponding to stronger attention.

**(B)** Magnified view of the representative attention map (layer 6, head 5) indicated by the red rectangle in panel **A**. The colour scale is identical to that shown in panel **A**.

**Supplementary Figure 3.** Training dynamics of context-aware models

Training and validation loss curves for models incorporating biological context.

**(A)** Training loss for the sequence + context model.

**(B)** Corresponding validation loss over epochs.

**(C)** Training loss for the sequence + context + motif features model.

**(D)** Corresponding validation loss over epochs.

**Supplementary Figure 4.** Training dynamics for gene-wise held-out models

Training loss curves for the remaining gene-wise held-out models (16 genes). Most models show consistent convergence behavior, with the exception of MED4 and CG10915, which display reduced predictive performance (R² < 0.4).

