## Supplementary figures and images for "Decoding Promoter Activity from DNA Sequence using Pre-trained Language Models"

### Supp. figs 1-4

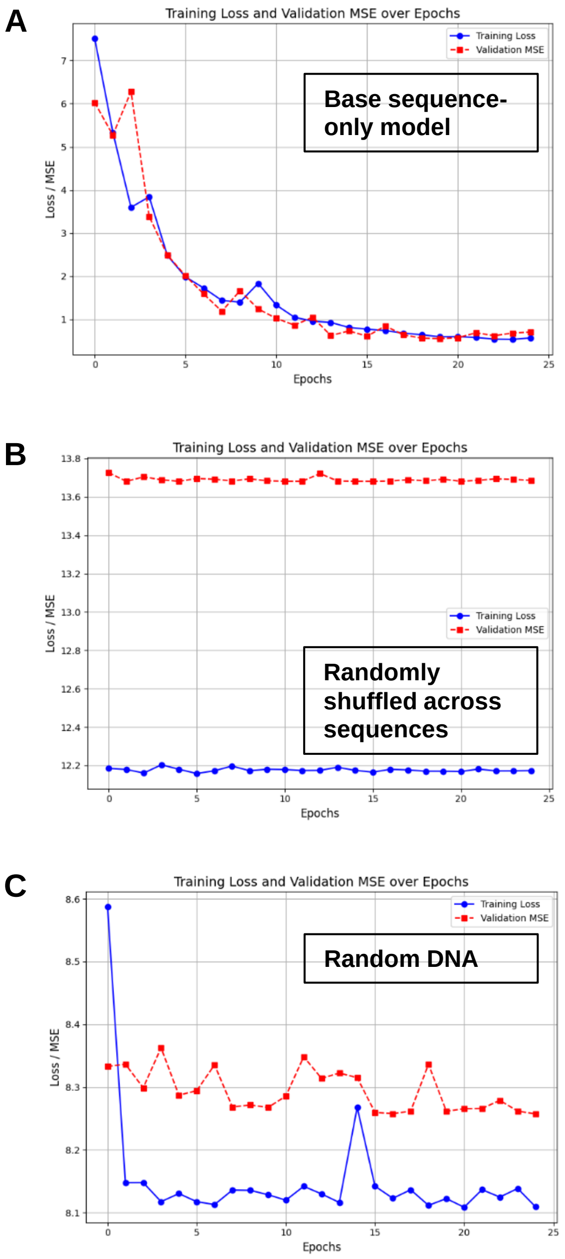


#### Supplementary Figure 1

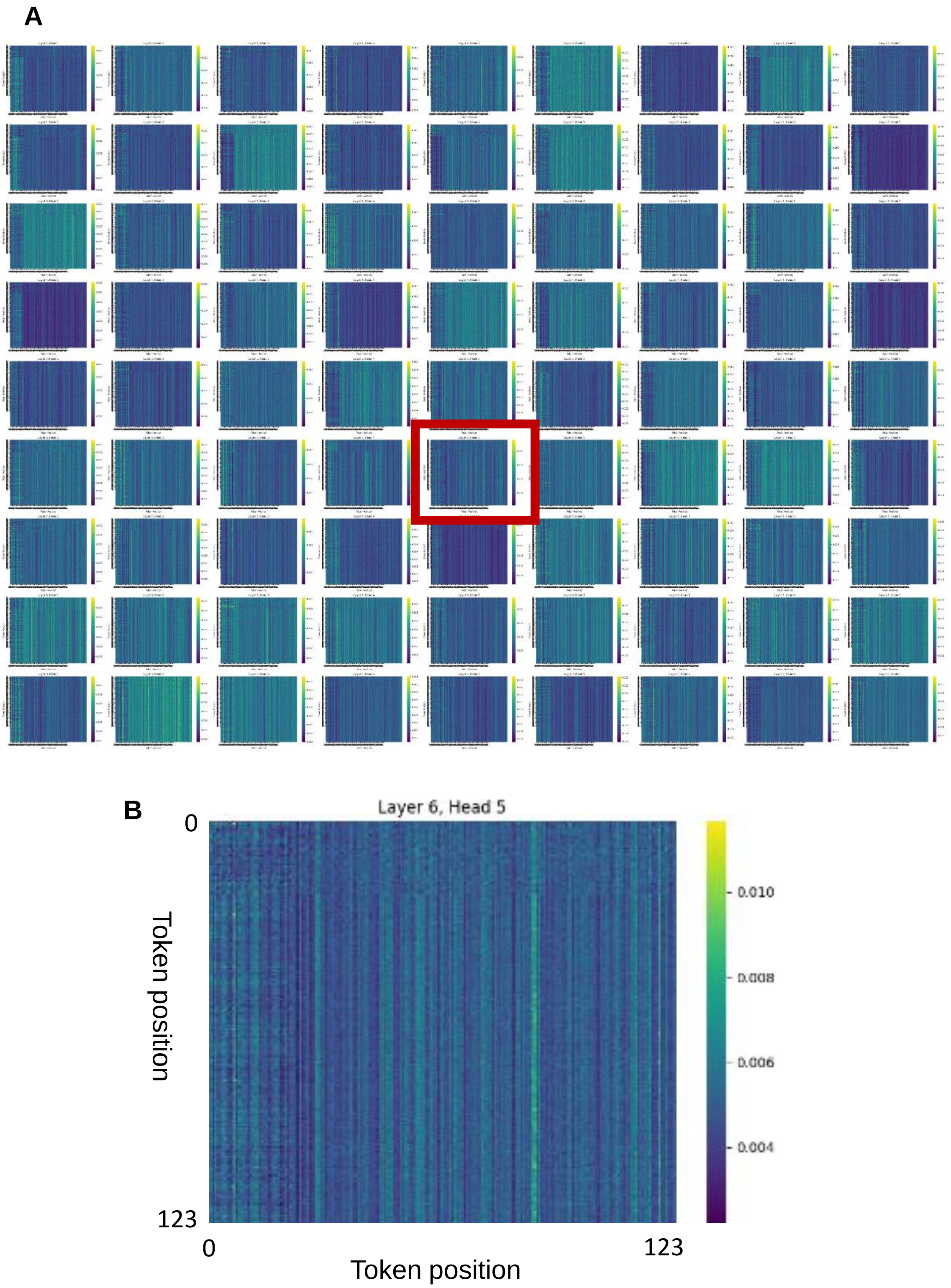


#### Supplementary Figure 2

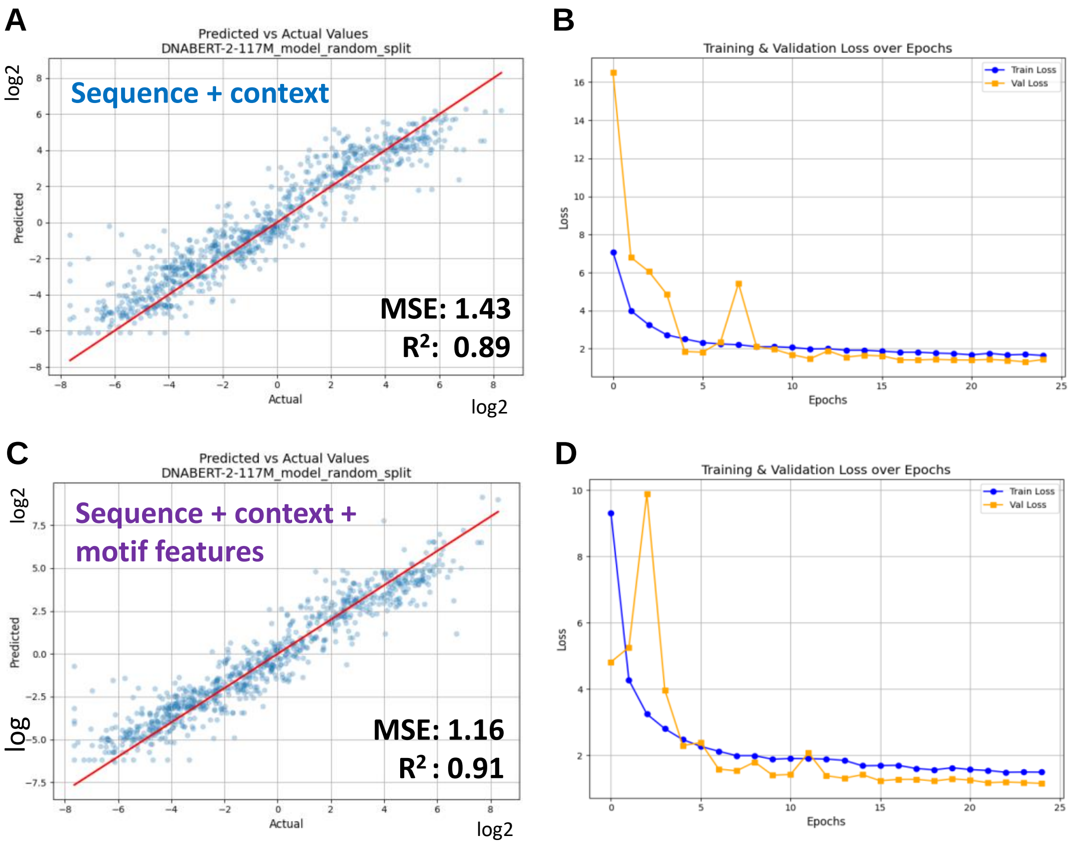


#### Supplementary Figure 3

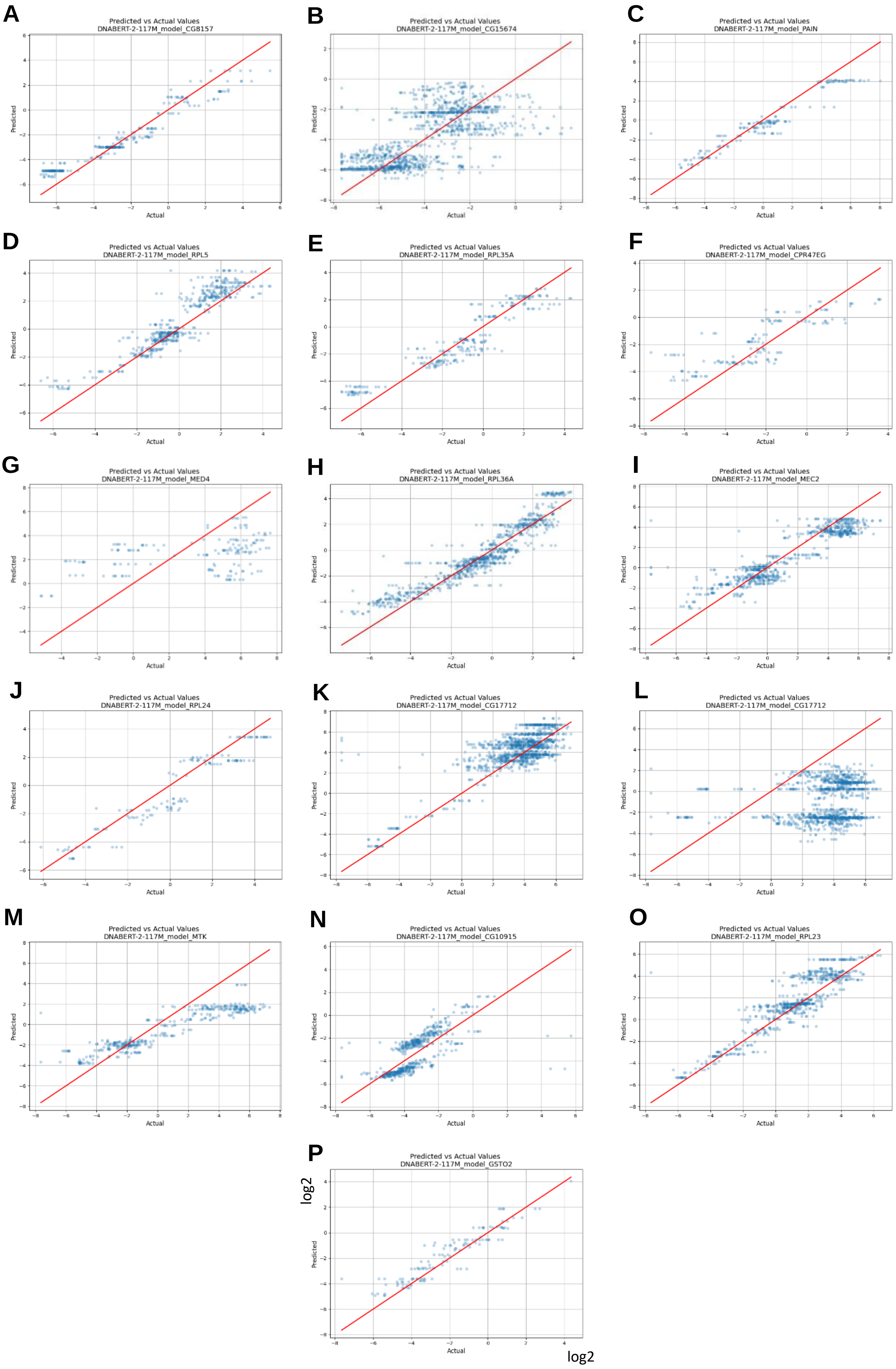


#### Supplementary Figure 4
